# SynDISCO: A Mechanistic Modeling-Based Framework for Predictive Prioritization of Synergistic Drug Combinations Directed at Cell Signaling Networks

**DOI:** 10.1101/2023.04.07.536087

**Authors:** Sung-Young Shin, Lan K. Nguyen

**Author notes:** To whom correspondence should be addressed (L.K.N) and (S-Y.S.).

## Abstract

The widespread development of resistance to cancer monotherapies has prompted the need to identify combinatorial treatment approaches that circumvent drug resistance and achieve more durable clinical benefit. However, given the vast space of possible combinations of existing drugs, the inaccessibility of drug screens to candidate targets with no available drugs, and the significant heterogeneity of cancers, exhaustive experimental testing of combination treatments remains highly impractical. There is thus an urgent need to develop computational approaches that complement experimental efforts and aid the identification and prioritization of effective drug combinations. Here, we provide a practical guide to SynDISCO, a computational framework that leverages mechanistic ODE modeling to predict and prioritize synergistic combination treatments directed at signaling networks. We demonstrate the key steps of SynDISCO and its application to the EGFR-MET signaling network in triple negative breast cancer as an illustrative example. SynDISCO is, however, a network- and cancer-independent framework, and given a suitable ODE model of the network of interest, it could be leveraged to discover cancer-specific combination treatments.

## 1. Introduction

The inevitable development of resistance to anticancer monotherapies has highlighted the need to identify combinations of drug agents as a more effective approach to cancer treatment (1, 2). Indeed, combination therapies are being actively explored across all cancer indications (3). Yet, given the vast space of possible target combinations, we are faced with daunting challenges of how to predict and prioritize the best possible combinations of targets (and drugs targeting them) for clinical validation. Although experimental screenings of combinations of existing drug agents are becoming increasingly feasible (3), the large number of available drugs and the marked heterogeneity of cancers render exhaustive experimental screens costly and impractical. Moreover, experimental screens can only incorporate molecular targets that already have existing drug(s) but cannot examine candidate targets for which no drugs are available. Given the still rather sizable number of the latter, this presents a significant missed opportunity. There is therefore an urgent need to develop new computational approaches capable of interrogating a large number of drug combinations efficiently in an unbiased manner, from which their anti-tumor effects can be systematically predicted and prioritized.

Signaling pathways play a critical role in the governing of cancer-relevant cellular processes, including cell growth, death, migration and differentiation (4, 5). Reflecting this, signaling protein kinases represent a major class of cancer therapeutic targets. Following the FDA approval of the first kinase inhibitor imatinib in 2001, 20 years later in 2021, 62 therapeutic agents targeting about two dozen different protein kinases have been approved by the FDA, most of which are prescribed for the treatment of neoplasms (6). However, the number of kinases having approved drugs is dwarfed compared to the total number of kinases encoded in the human proteome (518 (6)), many of which have been implicated in tumorigenesis, leaving significant room for further therapeutic discovery.

However, the translation of kinases from promising candidates to clinically actionable targets has been slower than expected. Part of the reason is due to the enormous complexity of cellular signaling networks. These networks usually contain intricate post-translational modifications and complex intertwined feedback loops and elaborate pathway crosstalk, which together render it highly challenging to analyze and predict cellular response to drug treatment (7, 8). These complexities also complicate the search for potential synergistic therapeutic targets within signaling networks. To address these issues, mechanistic and dynamic mathematical models, such as Boolean network models (9, 10), dynamic Bayesian networks (11), fuzzy-logic models (12), and ordinary differential equation (ODE) network models (5, 13-16) have been developed for a variety of signaling networks. In addition to translating complex biochemical relations into precise mathematical terms, these models also provide powerful quantitative platforms for analyzing and predicting drug response, as well as inferring causal mechanisms that underscore nontrivial behaviors, all of which are critical in facilitating the identification of novel and effective drug combinations.

In this chapter, we present a practical guide to SynDISCO, a computational framework that leverages mechanistic ODE modeling to predict and prioritize synergistic combination treatments directed at signaling networks. We first demonstrate the key modules and steps of SynDISCO and then illustrate its application through a case study involving the EGFR-MET crosstalk network in triple negative breast cancer. SynDISCO is, however, independent of the signaling network or cancer type, and as long as an appropriate ODE model of the network is available, it could be integrated on top of the model to discover cancer-specific combination treatments.

## 2. Description of SynDISCO

**Figure 1.**
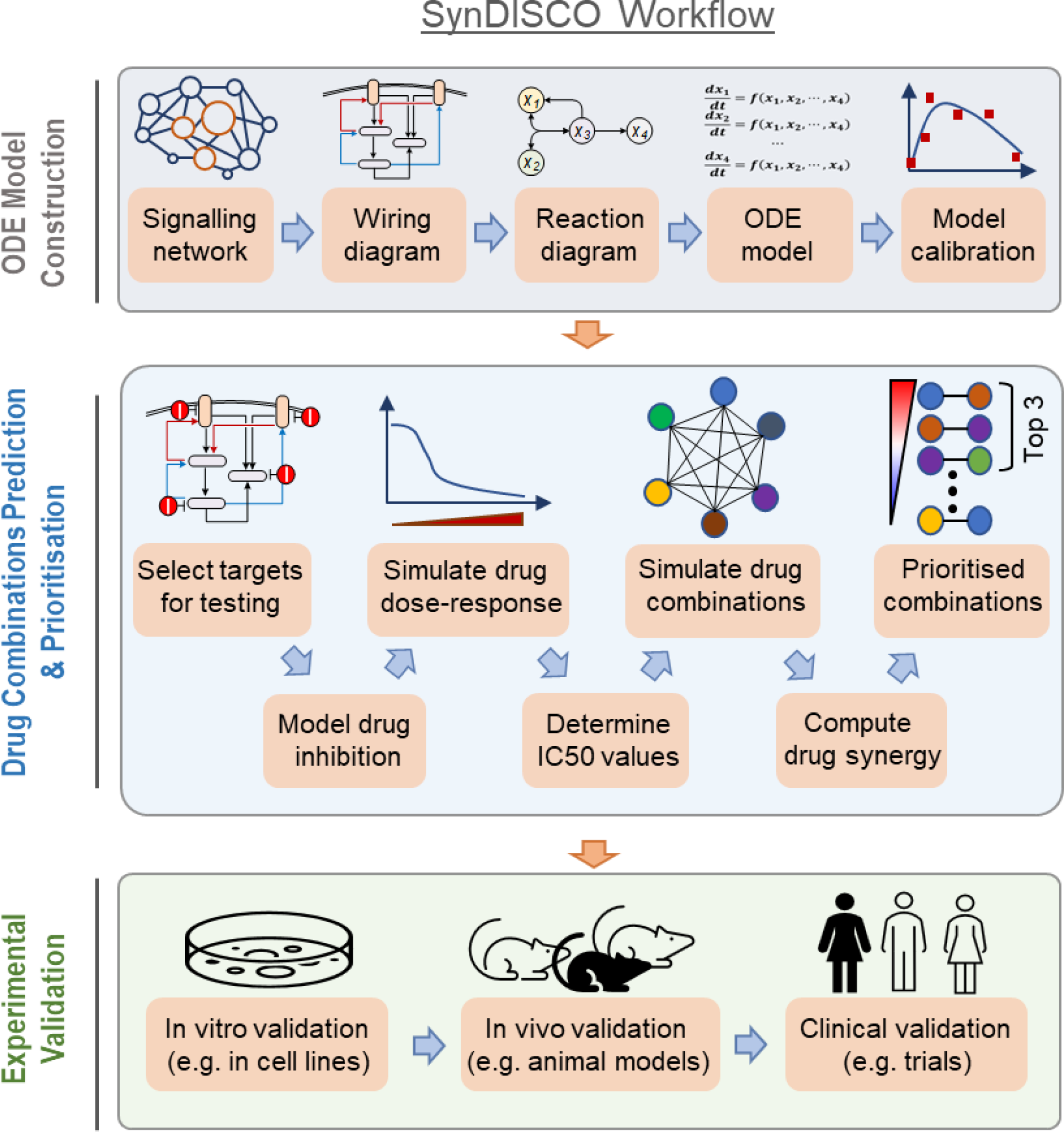
A workflow of SynDISCO. The pipeline contains three modules: ODE model construction (module 1), drug combinations prediction and prioritization (module 2) and experimental validation (module 3)

The entire SynDISCO pipeline consists of three modules, depicted in Figure 1. The first module involves construction of a predictive mathematical model in ODE format that quantitatively describes a signaling network/pathway of interest. This module, however, can be skipped if such a suitable model already exists. Building upon the ODE model from module 1, the second module involves computational simulation and evaluation of a specified number of pair-wise drug combinations directed at the network nodes (model species), both in terms of inhibitory effect and drug synergy. The drug combinations are then ranked and prioritiszd according to the levels of computed synergistic effect. Once the combinations are ranked, the top candidates will be tested experimentally, and the results will be cross-validated with model predictions in the last module, module 3. While it is highly desirable for module 3 to take place, in practice this is optional depending on the experimental capabilities of the investigator team. In such a case, the end result of SynDISCO would be a set of predicted promising drug combinations that can be followed up experimentally by other researchers. Below, we describe the steps within each module in more details, with an emphasis on module 2 due to its central importance within the whole pipeline.

### 2.1. Module 1: ODE Model Construction

Once a signaling network of interest is settled, construction of an ODE network model can begin. This typically involves stepwise establishment of a network wiring diagram, followed by a reaction schematic diagram, then formulation of differential equations from the model reactions, and finally by model calibration against known experimental data (17, 18).

Building a biologically accurate and evidence-based wiring diagram that specifies the network components and how they interact is the first and critical step in kinetic model development. A good starting point for this would be to consult signaling pathway databases (19), for example Reactomes (20), KEGG (21), Signor (22), or WikiPathways (23). However, since the wiring of the same signaling network may differ subtly between different tumor types (or even cell lines of the same tumor) while these databases typically do not contain context-specific interactions, one often has to refine the diagram using knowledge of interactions specific to the biological context of interest. These could come from in-house data, up-to-date reviews, or previous models developed for the same network (e.g., from Biomodels (24)). More often than not, conflicting information about specific links may arise, at which point one has to try to clarify and resolve, for instance through obtaining additional experimental data. In the case this is not possible and assumptions have to be made, they should be clearly stated.

Upon completion of the wiring diagram, it should be made further precise by converting it into a reaction schematic diagram (5, 13). Here, the interactions between network nodes are described by biochemical reactions with clear specification of the substrates, products, and catalyzing enzyme species. As an example, a positive regulatory link between a kinase (K) and its substrate (S) in the wiring diagram would be represented in the reaction diagram by a reaction: S → phosphorylated S (pS), catalyzed by K. The advantage of this step is that once it has been done, the ODEs can be straightforwardly translated from the reaction diagram without any ambiguity. In other words, the reaction diagram provides a one-to-one visual mapping to the ODEs that is convenient for model checking, modification and troubleshooting. Next, the rate equations for each reaction are constructed based on kinetic laws, for example, mass-action kinetics for association/ dissociation of molecules, Michaelis-Menten kinetics for (de)phosphorylation and (de)ubiquitination), and Hill kinetics for transcriptional regulation (17). A set of ODEs can then be built based on the rate equations and by considering appropriate production and elimination terms for each node.

The final step in module 1 is to calibrate the ODE model against known experimental data, which is critical in conferring the model with identity and predictive power. Building a model without this step is a job half done. Model calibration involves computational estimation of unknown model parameters using optimization algorithms with the goal to minimize the discrepancies between model simulation and experimental data. A variety of techniques are available for model parameter estimation, and we refer readers to an excellent recent review by Villaverde et al. (25) for detailed information regarding what best to and not to do. The ideal end product here is a well-calibrated ODE model of the signaling network under study.

### 2.2. Predictive Prioritization of Drug Combinations

Using the predictive ODE model from module 1 as input, module 2 aims to predict and prioritize various pair-wise drug combinations targeting the network nodes based on the levels of synergistic effect displayed by them. This module involves multiple steps described below.

#### Selection of Targets for Virtual Screening

Prior to simulation, a set of drug targets need to be specified. This involves specification of the pairs of targets, that is, suitable nodes within the modelled network that will be co-inhibited. It is important to note that the choice of targets for computational evaluation does not necessarily require them to have actual existing drugs, meaning suitable targets without any targeting drugs can be considered (see **Note 1**).

#### Modeling Drug Inhibition of Target Nodes

Following the selection of target nodes, we assume that each of them in theory can be inhibited by a corresponding small-molecule inhibitor. Hence, hereafter we will refer to target combinations and drug combinations interchangeably. The action of the inhibitor on its target needs to be explicitly incorporated into the existing ODE equations by modeling the target-inhibitor interactions.

In this regard, two common modes of target inhibition can be considered: competitive and allosteric inhibition. Consider a simple catalytic reaction where a kinase binds the substrate (S) to form a complex (ES) before producing the product (P) (Fig. 2). In competitive inhibition, an inhibitor (I) binds to the kinase and form a competitive, inhibitory complex (EI) (Fig. 2a). This type of inhibition can be completely overcome by very high concentration of the substrate S. Thus, the rate of the kinase-catalyzed reaction becomes saturated at a certain point (same *V_m_*), whereas the Michaelis constant (*K_m_*) is increased by a factor of (1 + [I]/K_i_), where [I] is the drug concentration and K_i_ is the dissociation constant (Fig. 2c).

On the other hand, in allosteric inhibition, the kinase has an additional binding site other than the site for the substrate (26), where an inhibitor binds, forming both a binary (ES) and ternary complex (ESI) (Fig. 2b). Unlike competitive inhibition, in allosteric inhibition *V_m_* changes by a factor of (1 + [I]/K_i_) while *K_m_* is not affected (Fig. 2b, d).

**Figure 2.**
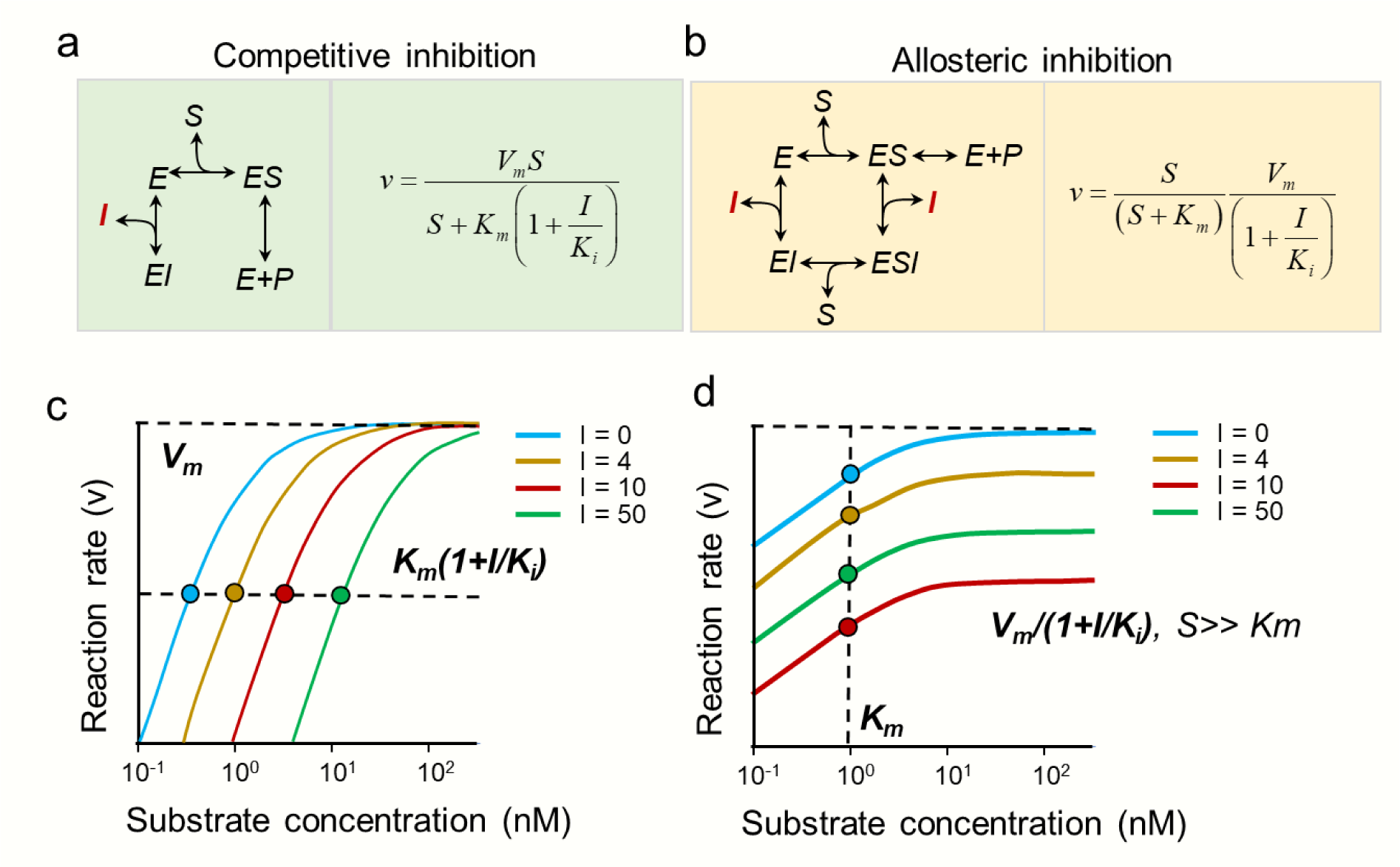
**Mathematical formulation of different modes of drug-target interaction**. (a) Competitive inhibition. (b) Allosteric inhibition. E, enzyme; S, substrate; I, inhibitor; P, product; (c-d) Substrate–rate curve for competitive inhibition (c) and allosteric inhibition (d)

#### Simulating Drug Dose-Response Curves

Having incorporated the actions of the inhibitors, one can simulate the effect of drug treatment under various conditions. It is important to make sure the model is under the steady state before application of the drugs (Fig. 3a), in order to better mimic experimental scenarios (where cells are often grown in cultured medium before drug treatment). However, in general specific model simulations should be guided by the exact experimental setup (*see* **Note 2**). Next, to determine the dynamic responses to a range of drug concentrations for subsequent analyzes, dose-response simulation should be performed (Fig. 3b). The most obvious response readout would be the activity of the corresponding drug target but can also be a (theoretical) function representing cell growth or viability (*see* **Note 3**). Dose-response curves are established by gradually increasing the concentration of the drugs and simulating the levels of the response readouts, often at a time point where steady state is reached after drug treatment.

**Figure 3.**
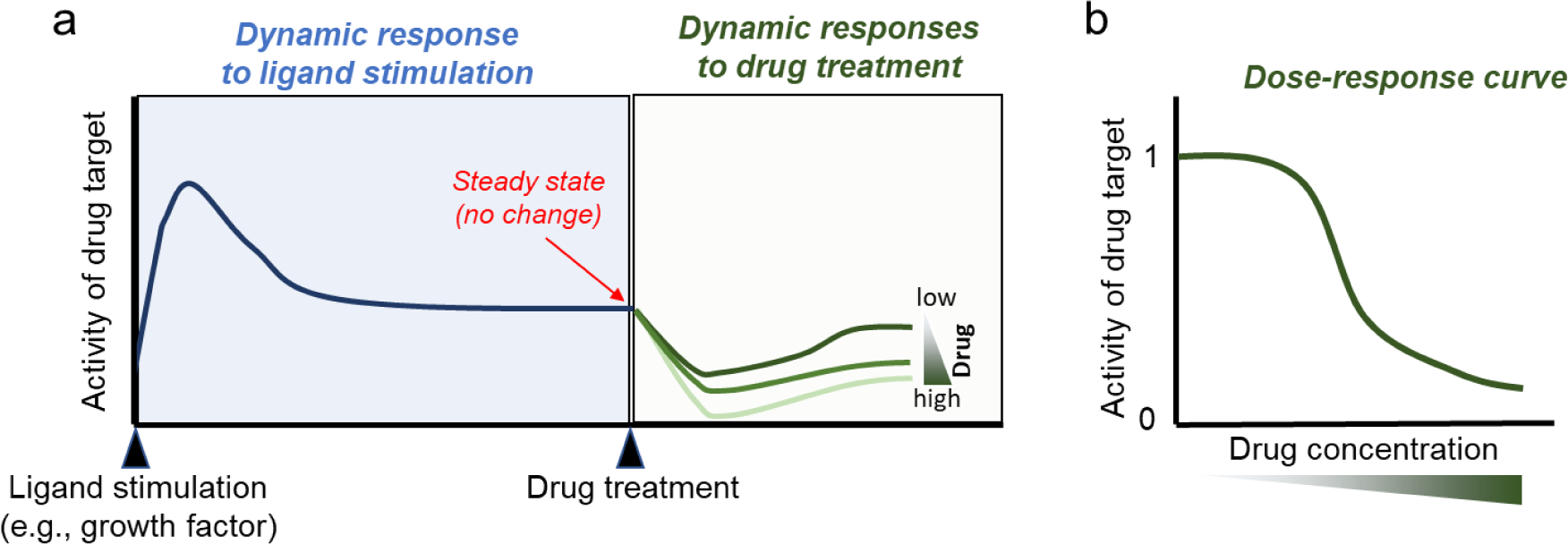
An illustration of a time-dependent drug response simulation. It consists of two-step simulations: ligand stimulation and drug treatment (a). Dose-response curve measured at a specific time point (e.g., 24 h after a drug treatment) (b)

#### Determination of Drugs’ IC50 Values

The half-maximal inhibitory concentration (IC50) is a commonly used value for quantifying the potency of a drug and is often employed in drug combination studies. The IC50 is also used as a reference value of a drug concentration because drugs have different sensitivity for different cells. Thus, in an experimental design, one uses a fold of IC50 concentration (e.g., IC25 or IC50) to treat cells rather using absolute drug concentration. Here, the theoretical IC50 value for each drug inhibitor will be computed based on the dose-response curves from the previous step by fitting these curves to the four-parameter dose-response function (27) depicted in Figure 4.

**Figure 4.**
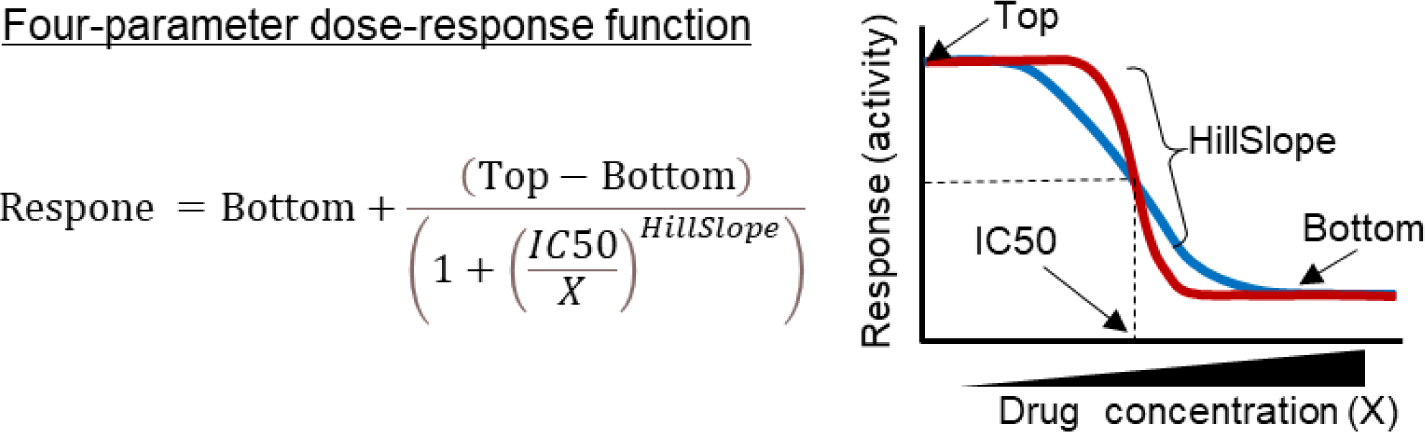
The four-parameter dose-response model for estimation of IC50. Top is the maximal value of response curve and Bottom is the minimal value (i.e., maximally inhibited response). Top and Bottom are plateaus of the curve in the units of the Y axis. HillSlope denotes the steepness of the sigmoidal curve. When HillSlope = 1, the dose response is “standard”; when HillSlope <1, the curve becomes “shallower”; when HillSlope >1, the curve becomes “steeper”

#### Simulating the Effect of Drug Pairs

Having determined the suitable dose for each drug (e.g., IC50), here the effect of the pair-wise drug combinations can be systematically simulated and compared. Because in most scenarios we want to inhibit the growth/viability of the cancer cells, thus the effect of the drug combinations on a molecular readout that best reflect cell growth/viability can be considered. Alternatively, as discussed above and in **Note 3**, one can use a heuristic function representing cell viability as the readout to judge the effect of drug treatment. In any cases, for each drug combination consisting of drug A and B, the effects of vehicle treatment (no drug), drug A alone, drug B alone, and their combination are simulated.

A key way in which SynDISCO assesses drug combinations is to quantify possible synergy (or lack thereof) between the individual drugs. To this end, different approaches can be used to compute drug synergism, depending on the specific context and preference. These include the Coefficient of Drug Interaction (CDI) model (28, 29), the Bliss Independence (BI) model (30, 31), the Chou-Talalay model (32), highest single agent (HSA) model (33) and zero interaction potency (ZIP) model (34, 35). More details about these techniques are given in **Notes 4** and **5**. Regardless of the chosen approach, a synergy score is derived for each drug combination. For example, using BI, based on the single-drug and combined drug effects previously simulated, a synergy score for each drug pair is computed where the synergy score <1, = 1 or >1 indicates that the drugs are synergistic, additive or antagonistic, respectively. Importantly, a smaller BI score indicates stronger synergism, while a larger BI score indicates stronger antagonism. Taking advantage of this quantitative property, SynDISCO then ranks all the tested combinations in order from most synergistic to most antagonistic, enabling prioritization of the combinations (Fig. 5).

**Figure 5.**
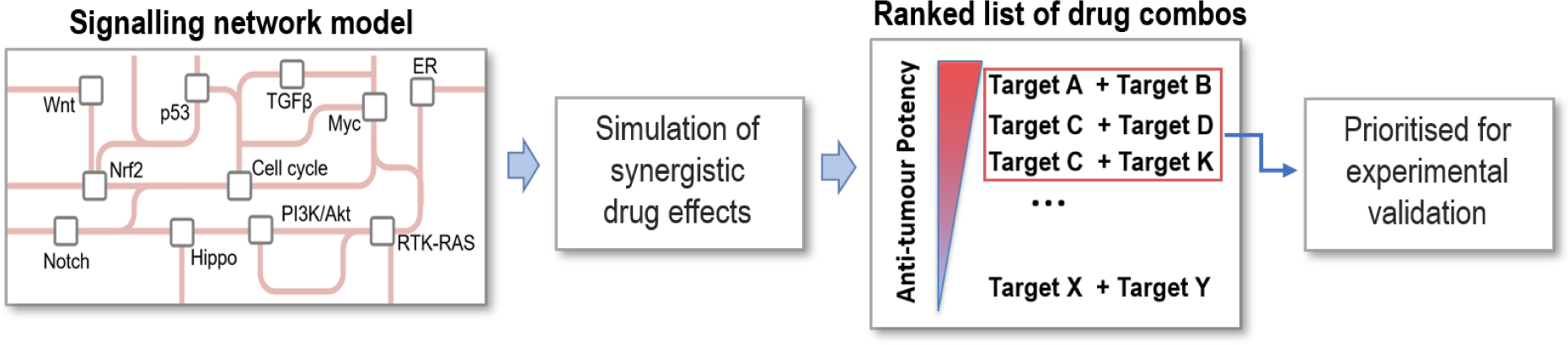
A workflow of identification and prioritization of combinatorial drugs

### 2.3. Experimental Validation of Top Candidates

By this stage, one should have obtained a set of highly synergistic drug combinations predicted by modeling that are ready for experimental validation. As mentioned earlier, although this module is optional depending on whether the investigators have the capacity to perform experiments, it is highly recommended as it is the only way to find out if the model predictions are correct or not. More importantly, any discrepancies between the experimental data and prediction will help pinpoint gaps in the model, allowing it to be further refined and improved.

One of the most common ways to test candidate cancer drug combinations is to perform experiments in cancer cell lines cultured in the laboratory. For example, the effect of single-drug and combination drug treatments, as well as vehicle control (e.g., DMSO) on cell viability or colony formation ability can be done using MTS/MTT assays and/or colony formation assay, respectively. The MTS assay is normally used to assess cell proliferation, cell viability, and cytotoxicity based on the reduction of the MTS (3-(4,5-dimethylthiazol-2-yl)-5-(3-carboxymethoxyphenyl)-2-(4-sulfophenyl)-2H-tetrazolium) compound by viable mammalian cells (36). The MTT (3-[4,5-dimethylthiazol-2-yl]-2,5 diphenyl tetrazolium bromide) assay is a colorimetric assay for assessing cell metabolic activity by measuring NAD(P)H-dependent cellular oxidoreductase, which reflects the number of viable cells present (37). On the other hands, colony formation assay (or clonogenic assay) is an in vitro cell survival assay based on the ability of a single cell to grow into a colony (38). Other methods such as the Incucyte or xCelligence systems can be also used to visualize and quantify living cell behavior over time by automatically gathering and analyzing images (39, 40). Promising candidates validated in cell lines can then be further tested in more advanced models such as organoids or mouse xenograft cancer models.

## 3. Application of SynDISCO to the EGFR-PYK2 Signaling Network in TNBC

In this section, we provide a concrete application by illustrating how SynDISCO is used to identify effective drug combinations directed at the EGFR-MET signaling network in triple negative breast cancer (TNBC), an aggressive subtype of breast cancer (13). The epidermal growth factor receptor (EGFR) and hepatocyte growth factor receptor (HGFR or MET) are frequently overexpressed in TNBC, making them promising therapeutic targets (41-43). However, single-drug inhibition of these kinase receptors failed to induce durable clinical benefit, primarily due to emergence of drug resistance (44, 45). Identification of more effective combinatorial therapies that circumvent resistance is therefore of urgent importance for TNBC.

### 3.1. Construction of the EGFR-MET Network Model

#### Building Reaction Diagram

Given our purpose here is to predict effective drug combinations targeting the EGFR-MET crosstalk signaling network in TNBC, our first step is thus to establish an ODE model of this network in the context of TNBC (Fig. 6a).

#### ODE Model Formation

From the reaction diagram, a set of ODEs was formulated using a combination of kinetic laws. Specifically, we employed Michaelis-Menten (MM) kinetics for (de)phosphorylation and (de)ubiquitination reactions; mass-action kinetics for association/dissociation reactions, and Hill kinetics for transcription reactions. In the end, the model consists of 13 ODEs and 25 reactions, which are given in Appendices A and B.

#### Model Calibration

As the last step of module 1, we calibrated the model using a dataset of kinetic, time-resolved data under different perturbation conditions, obtained from the TNBC cell line MDA-MB-468. This includes the time-dependent changes in the phosphorylated and total levels of key network components (EGFR, PYK2, MET, and STAT3) in response to EGF stimulation. A genetic algorithm-based global optimization procedure implemented in Matlab was used for model calibration and estimation of unknown model parameters (46, 47). As result of this process, a set of best-fitted parameter values was obtained (Appendix C). Model simulations using the best-fitted parameter set recapitulate the kinetic profiles of the observed data, as shown in Figure 6b. The calibrated model now can be used for the next steps in module 2.

**Figure 6.**
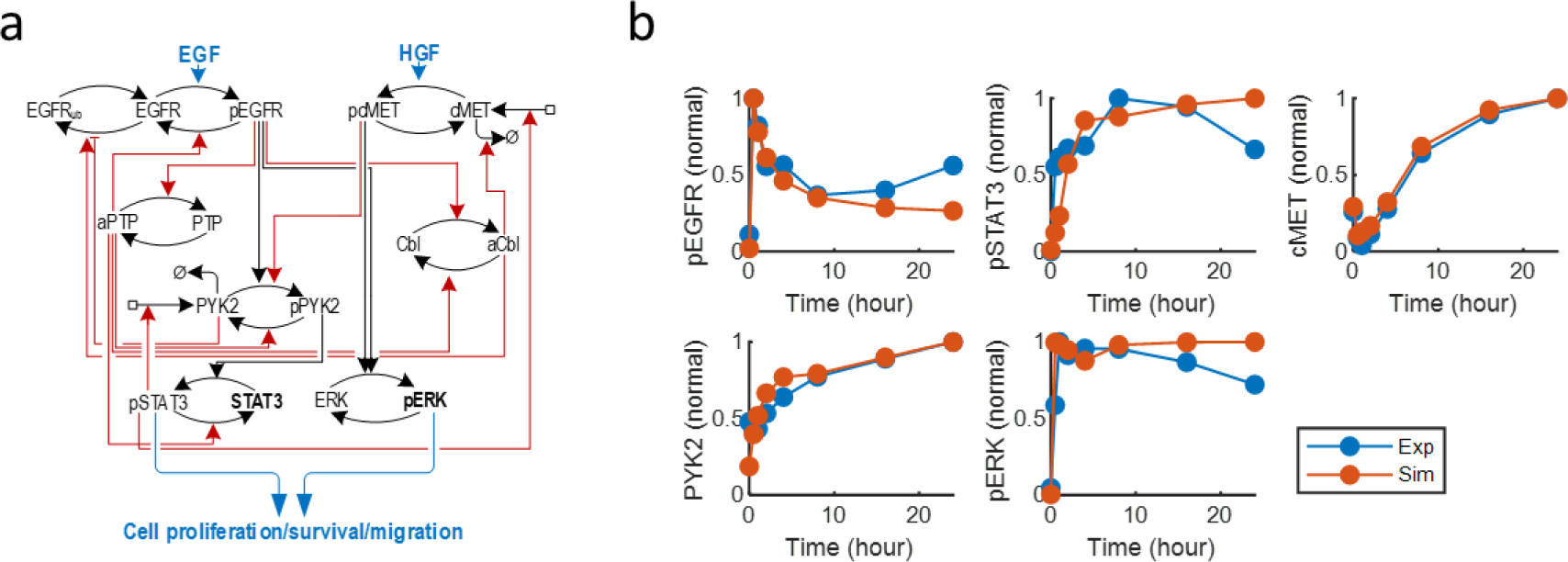
An EGFR-MET network model and model calibration (13). (A) A schematic diagram of EGFR-MET network model. (B) The model calibration and fitting to experimental data. Red line: simulated data of the best-fitted model. Blue line: experimentally observed time-course data

### 3.2. Drug Response Simulation of the EGFR-MET Model

#### Selecting Targets

We select EGFR, PYK2, STAT3 and MET as possible targets for exploration because they are known pro-growth kinases, and are frequently altered in TNBC (13). About half of TNBC overexpress EGFR (48). Proteomics analysis revealed that PYK2 is active mainly in basal-like breast carcinomas among breast cancer subtypes, and activated PYK2 correlates with high levels of EGFR (42). Moreover, STAT3 frequently is activated in TNBC cells (49), while MET is highly expressed in ∼52% of TNBC (50).

#### Modeling Drug Inhibition

We assumed in the model that the target EGFR, PYK2, MET and STAT3 can be inhibited by their respective small-molecular inhibitors: Gefitinib, PF396, EMD-1214063 (EMD) and Stattic, respectively. We assumed Gefitinib effectively blocks the phosphorylation of major EGFR tyrosine sites (51). PF396 inhibits the kinase activity of PYK2 by preventing the transfer of a phosphate group to a target protein from ATP (52). EMD is a small molecule that inhibits c-Met phosphorylation of target substrates (53, 54). In particular, we assumed that Gefitinib, PF396 and EMD allosterically inhibit their targets since there is no evidence that they compete for the binding sites with the activating kinases (13). On the other hand, Stattic was modelled as a mass action interaction with a target molecule due to its a reversible drug-target binding reaction (55).

#### Simulating Drug Dose-Response and Determining IC50 Values

Here, we assumed the phosphorylated levels of ERK and STAT3 (pERK and pSTAT3), the two major oncogenic signaling effectors downstream of EGFR and MET as the models’ co-outputs that are used as surrogate markers of cell growth in order to assess the effect of drug treatments. Drug potency is therefore determined by the degree to which the drug inhibits these readouts.

As an example, Figure 7a displays the simulated dose-response curves for each drug, using pSTAT3 as a readout. These curves were then fitted to the Hill-type function discussed in Sect. 2. The estimated values of IC50 along with other function parameters for each drug are displayed in Figure 7b. The estimated IC50 values provide the drug-specific doses as basis for simulation of drug combinations in the next section. Similar steps should also be done for pERK as the readout in order to obtain drug specific doses specific for this readout (not shown).

**Figure 7.**
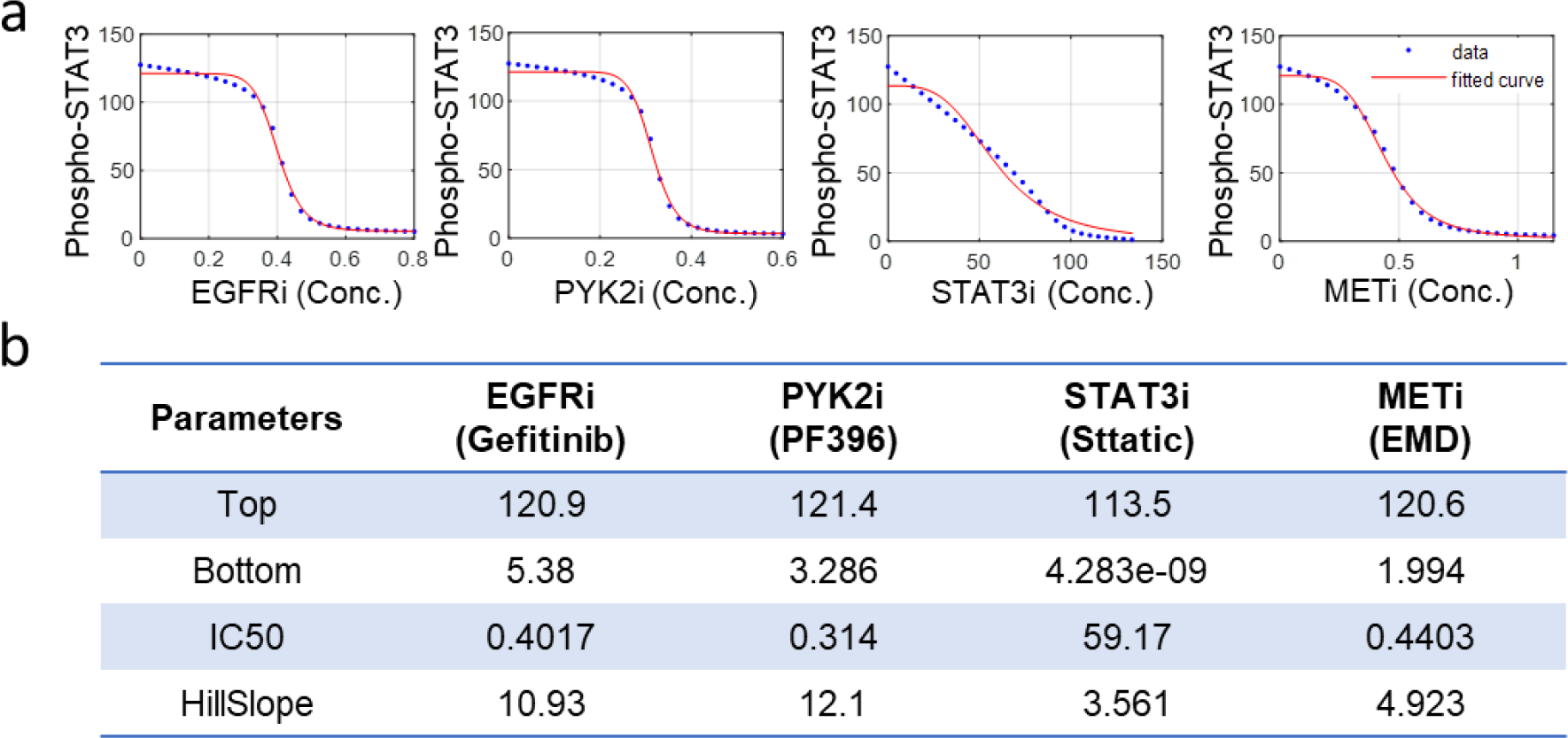
Estimation of IC50 using the four-parameter dose-response model. (a) Dose-response curves of pSTAT3. (b) The estimated parameter values of the dose-response function

#### Simulating Drug Combination Effect and Drug Synergy

With four drugs, we have a total of six different pair-wise combinations for evaluation. For each drug pair, we simulated the steady-state response of the readouts, that is, pERK and pSTAT3, to the single-drug treatments, the combined treatment at the pre-determined dose (e.g., IC25 or IC50) of each drug, and vehicle (no drugs) treatment. The drug synergy score for each drug pair can then be computed at this dose combination, using different synergy scoring methods such as CDI or BI (*see* **Note 4**). Figure 8a displays the computed synergy scores for the six drug combinations using pSTAT3 as readout, ordered from the most synergistic to most antagonistic combinations based on the values of the score. This ranked list allowed us to identify the most synergistic combinations which will be prioritiszd for experimental validation. Specifically, the model predicted that among the six combinations, dual inhibition of EGFR-PYK2 (using Gefitinib + PF396) and EGFR-MET (using Gefitinib + EMD) represents the most synergistic combinations, a prediction consistently made by both CDI and BI (Fig. 8a). Importantly, we also obtained similar ranking of the combinations when pERK was used as the readout (Fig. 8b). This led us to hypothesize that co-targeting EGFR-PYK2 or EGFR-c-Met will yield synergistic effects in inhibiting TNBC growth.

The concordance in terms of drug combination ranking between the readouts makes it easier to prioritize the top combinations, however it is possible that the predicted ranking may differ between different signaling readouts. In this case, one may consider using a single composite function of multiple signaling readouts as an indicator of cell viability (*see* **Note 3**).

Once highly synergistic drug combinations are predicted, further analyzes are recommended to obtain deeper insights into the mechanistic basis of how synergy is gained. For example, one can interrogate the time-course response of the network in single vs. combined drug treatments. A common reason for synergy is that single-drug treatments tend to trigger unwanted rebound in oncogenic activity of specific network nodes, and combined drug treatments can eliminate such rebound and enable more durable blockade of oncogenic signals (2, 8).

**Figure 8.**
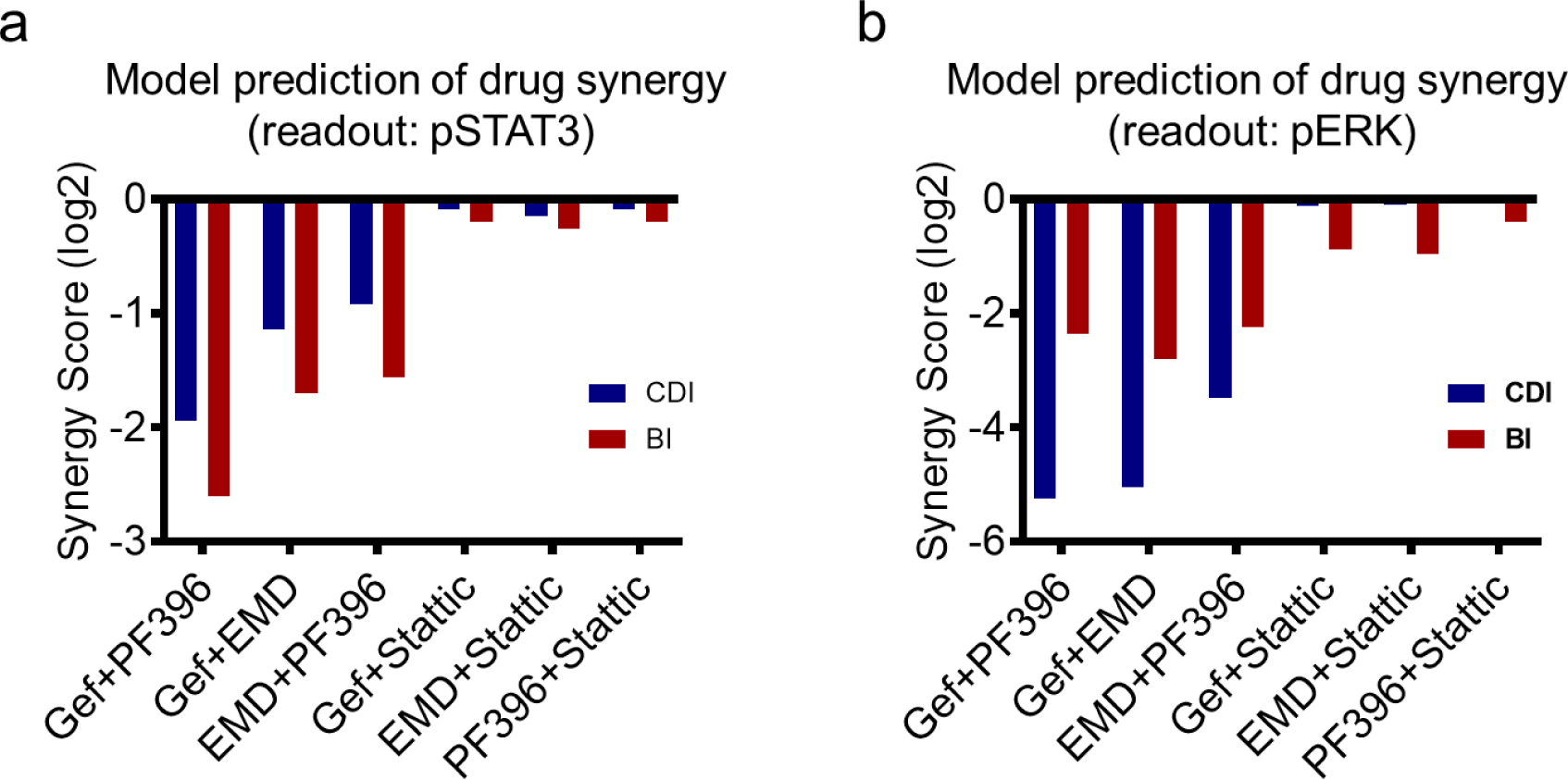
**Ranking synergy scores of drug combinations**. pSTAT3 (a) and pERK (b) were used as a readout for calculation of the synergy score. IC25 concentrations were used for each drug (note that similar results were found using IC50 concentrations)

#### Simulation at Varying Dose Combinations

The simulations above have primarily focused on testing the drug combinations by combining the drugs at specific doses (e.g. IC50 or IC25). However, a strength of our model-based approach is that it allows one to simulate drug combinations at myriad dose combinations within specified dose ranges, thus enabling a broader view of combinatorial drug effects. Using the top-predicted EGFR-PYK2 combination as an illustrative example, Figure 9a displays the simulated effects of dual inhibition of EGFR and PYK2 on the levels of pSTAT3 at increasing combined doses of the individual drugs. Follow on from this, Figure 9b displays the landscape of synergy score (CDI) computed at the corresponding dose combinations. These simulations revealed that the synergy score can vary significantly depending on the combined drug doses. Co-targeting EGFR-PYK2 displayed strongest synergism for the CDI score at around the IC50 of the drugs and for the BI score about IC25 (Fig. 9c) but exhibited antagonism at high drug concentrations. Although the synergy score landscape differs slightly between different drug synergy models because of different model assumptions, the above observation was supported by all the models. It is a good practice to perform different drug synergy models for cross-comparison (*see* **Note 5)**.

**Figure 9.**
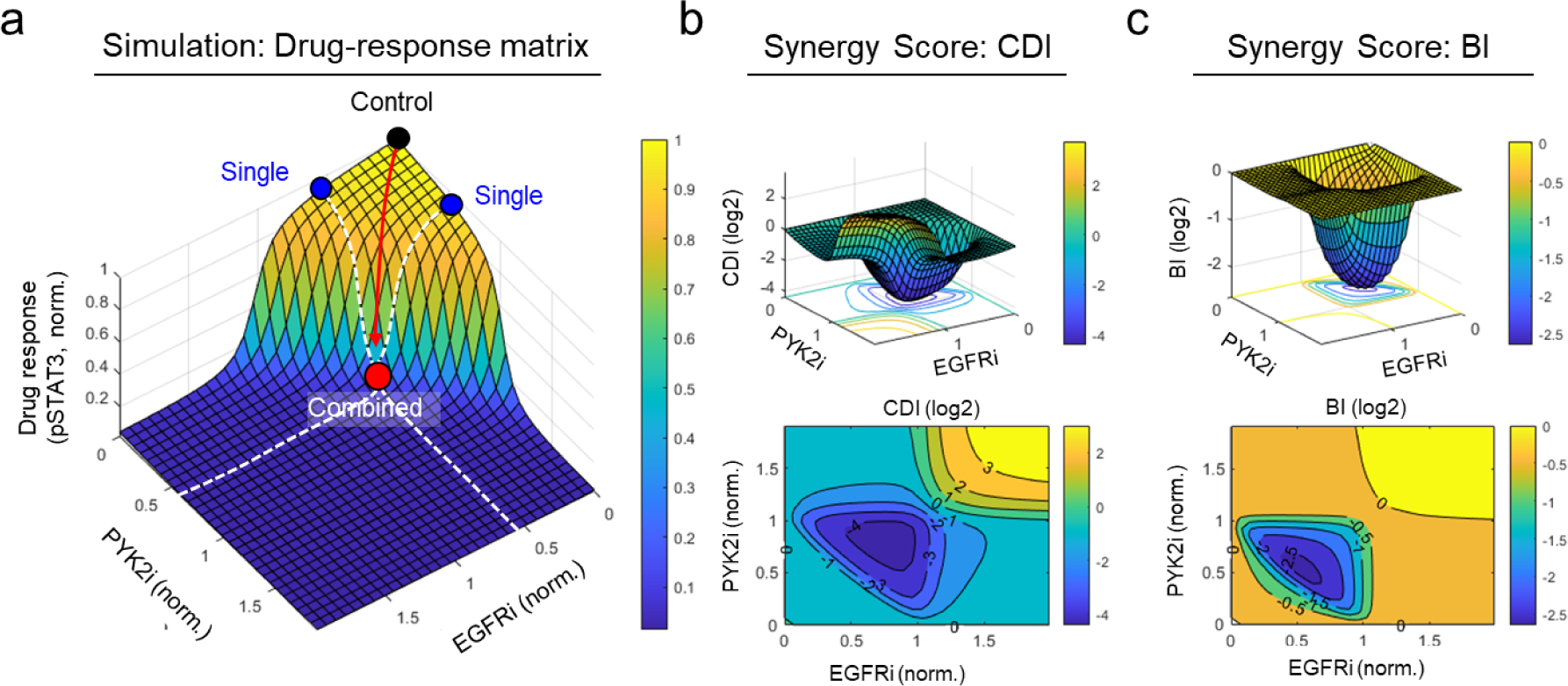
Simulation of pair-wise drug combination and computing drug synergism. (a) Drug response of pSTAT3 to EGFR and PYK2 inhibitors. Activation state of STAT3 was slightly suppressed upon single-drug treatments (blue dots), while the combined treatment significantly suppressed these activation states (red dot). (b) Synergy score of CDI (b) and BI (c). Note the drug concentration was normalized to the IC50 value of each drug. “1” indicates the IC50 value.

### 3.3 Experimental Validation of Top-Predicted Drug Combinations

It is desirable to verify model predictions with follow-up experiments. To experimentally validate the predictions in module 2, TNBC cells MDA-MB-468 were subject to increasing levels of the inhibitors Gefitinib, PF396, EMD, and Stattic in six pair-wise combination schemes, and the effects of the single- and combined-drug treatments on cell viability were measured using the MTT assay (13). As an example, Figure 10a presents the data for co-targeting EGFR and PYK2 using Gefitinib + PF396 (*see* **Note 5** for more information on experimental setup of drug treatment).

To analyze the synergism of all six drug combinations, we calculated the synergy scores using the obtained data based on the Chou-Talalay method (Fig. 10b). Consistent with model prediction, the data-based ranking of the combinations indicated that dual blockade of EGFR-PYK2 using Gefitinib + PF396, respectively, was indeed the most synergistic combination, followed by EGFR-MET co-targeting by Gefitinib + EMD. In addition, the ranked order of the six drug combinations was largely in concordance with model prediction. Importantly, co-targeting of EGFR and PYK2 was also shown to significantly reduce the growth of basal-like TNBC cells in animal models, and it was found that this combination was more potent than dual inhibition of EGFR and Met (42). Together, these experimental studies support the model predictions.

**Figure 10.**
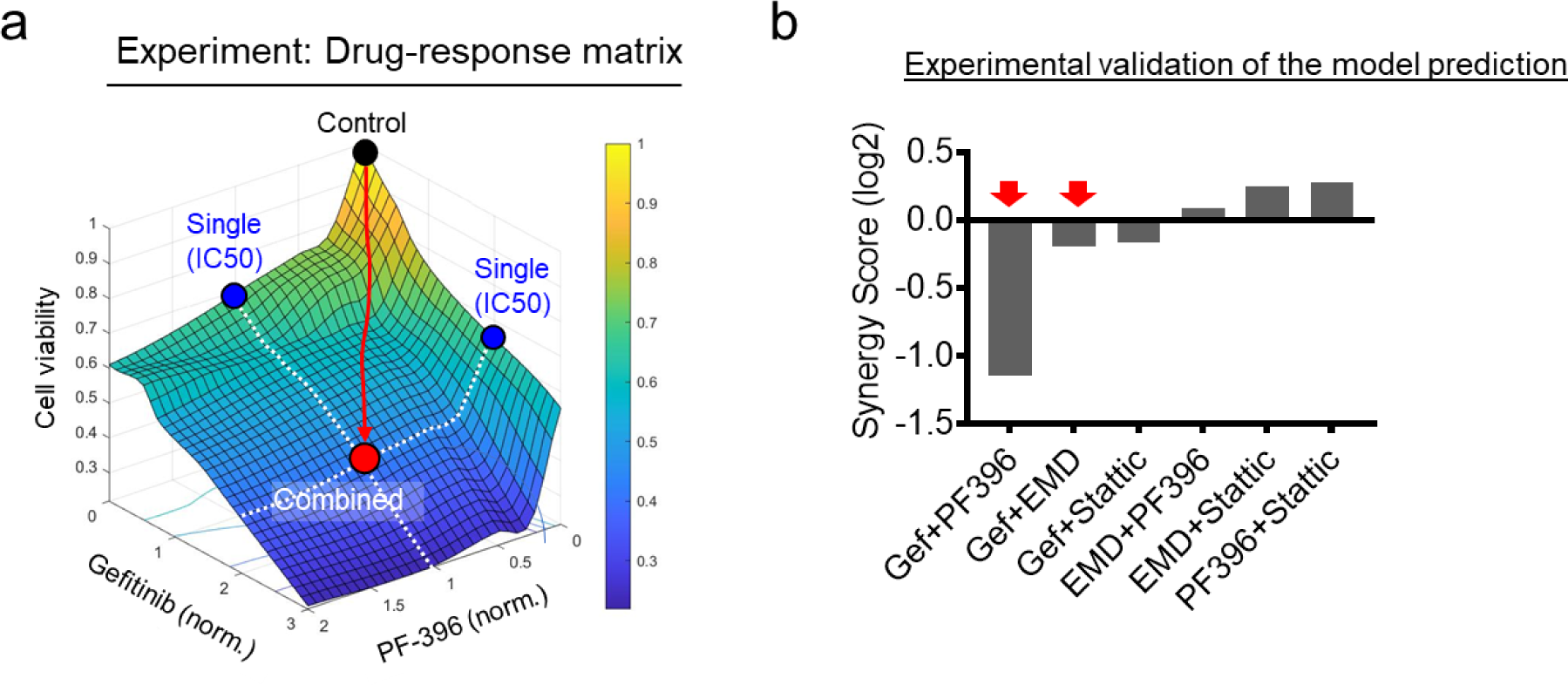
**Experimental validation of pair-wise drug combination**. (a) Drug response of cell viability to Gefitinib and PF-396. (b) Ranking synergy score of drug combinations. IC50 concentrations were used for each drug

## 4. Summary

Resistance to anticancer treatments, particularly single-agent therapies, remains a major clinical barrier to a cure in most cancer types. This highlights the critical need to predictively identify combination therapies of clinical potential in an unbiased, scalable, and cheap manner. These capabilities would enable the most promising candidates to be tested experimentally in a selective rather than brute-force strategy, thereby saving cost and accelerating the bench-to-bedside translation for cancer drugs. While computation modeling and simulation have been an integral part of the cancer systems biology field (5, 13, 16, 47), modeling-based approaches that predict synergistic drug combinations remain limited. In this chapter, we have provided a pedagogical guide to SynDISCO, a holistic framework developed precisely for this purpose. We have demonstrated the applicability of SynDISCO through identification of synergistic drug combinations against triple negative breast cancer. We hope it will serve as a useful tool for discovery of potentially effective drug combinations in other cancer settings.

## 5. Notes

### **1.** Selection of Potential Targets

While experimental screens for drug combinations are limited by the availability of the drugs, this is not the case for SynDISCO-based screening. In principle, any network nodes can be included in the pool of potential targets for combination exploration. If it turns out that a node (with no targeted drugs) is robustly predicted to be a promising co-target in specific combinations, this may serve as a motivation to develop new compounds against that target. This is useful because the complexity of signaling network wirings may mask the therapeutic vulnerabilities of certain network nodes, and their hidden therapeutic potential can be revealed by computational simulations. On the flip side, in SynDISCO, the exploration of drug targets is constrained by the scope and size of the input ODE models since one cannot investigate targets that are not included in the model. Thus, the plan regarding which targets to be explored should be considered early and factored into the model construction step in module 1 to determine the model scope.

While the EGFR-MET example above represents a proof-of-principle case study with a small number of drug combinations, this number can be scaled up significantly given a larger ODE model. In other studies, we have used SynDISCO to evaluate tens of targets and hundreds of possible combinations without issues (56). Generally, there is a trade-off between the number of network nodes and the computational cost required to evaluate all possible target/drug pairs, and one needs to consider this balance, taking into account available computing capabilities.

### **2.** Dynamic Simulations

The drug dose-response curves in module 2 can be generated by measuring the readouts (i.e., responses) at different time points post drug treatment. A good approach is to get a sense of how long it would take for the network to settle into steady state following drug perturbations by performing long-term time-course simulations. For example, if the system takes 24 h to reach steady state, this time point could be used for the dose-response simulations (Fig. 3b). Sometimes, one may be interested in interrogating the acute effect of the drugs, and much earlier time points (e.g., 1 h) could be used instead.

### **3.** Heuristic Function Representing Cell Viability

Because the outputs of ODE models of signaling networks are signaling and not phenotypic readouts (e.g. cell viability), the effects of drug treatment on phenotypic responses cannot be directly derived. Rather, these are usually indirectly determined. One approach is to assume key oncogenic downstream model outputs (e.g., pSTAT3 and pERK, as in Sect. 3) as proxies of cell phenotypic responses (often cell viability). However, in the case where cell viability is best represented by multiple model outputs and/or when there are significant inconsistencies in the results between different outputs, then a composite function of these outputs could be formulated. A simple approach is to assume that the selected model outputs contribute linearly to the cell viability function as follows (57):

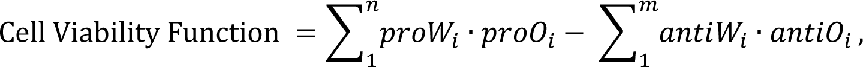

where *proO_i_* represents the *n* contributing pro-growth model outputs, while *antiO_i_* represents the *m* contributing anti-growth model outputs (e.g., proapoptotic nodes), and *proW_i_* and *antiW_i_* are the corresponding weights (normalized, between 0 and 1). Determining the weights of the outputs is a nontrivial task and may require further data and/or assumptions. For example, these weights can be estimated as part of the model calibration procedure given suitable experimental data that links cell viability to signaling readouts (57). In the absence of such data, a reasonable starting point may be to assume equivalent weights (i.e., equal 1).

### **4.** Methods to Compute Drug Synergy

There are a number of different methods for calculating drug synergy, underscored by different mathematical theories. These include the Coefficient of Drug Interaction (CDI) model (28, 29), the Bliss Independence (BI) model (30, 31), the Chou-Talalay model (32), highest single agent (HSA) model (33), and zero interaction potency (ZIP) model (34, 35). Here, we provide a brief description of the two common and simpler methods, CDI, BI, as a guide to the topic. However, we refer readers to the cited references for detailed technical explanation of each method.

#### Coefficient of Drug Interaction

CDI is a simple model to evaluate the inhibitory effect of the drug combination (28, 29). The synergy score, is calculated as follows:

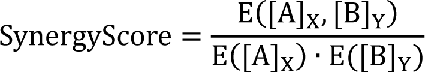

where E([A]_X_, [B]_Y_) denotes the combined drug response of drug A and B at X and Y concentration. E([A]_X_) and E([B]_Y_) are the single drug response at X and Y concentrations, respectively. Note the drug response should be normalized to its untreated control for the synergy calculation for using the BI model, and thus 1 indicate no drug treatment (ND) or control; 0 indicates the complete inhibition. SynergyScore <1, = 1 or >1 indicates that the drugs are synergistic, additive, or antagonistic, respectively. For instance, if compared to vehicle treatment (1), the tumor growth is decreased to 0.5 and 0.5 by the single-drug treatments, and to 0.1 by the combined treatment, then the SynergyScore = 0.1/(0.5 × 0.5) = is 0.4, indicating this drug combination has a synergistic effect (Fig. 11a).

#### Bliss Independence

BI is another statistical model to assess the combination efficacy of two drugs based on the assumption that the two drugs act through independent mechanisms and thus the individual drugs do not directly interfere with each other (30, 31). The predicted effect of drug A and B at drug concentrations of X and Y satisfies the following equation:

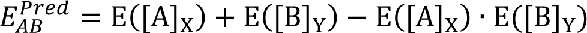

where 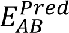 denotes the predicted combined effect of drug A and B at X and Y concentration. E([A]_X_) and E([B]_Y_) are the single drug effect at X and Y concentrations, respectively. Note that the drug response is indicated by a percent inhibition compared to an untreated control and thus 0 indicate no drug effect (0% inhibition) or control; 1 indicates the complete inhibition (100%). The BI score is calculated as follows:

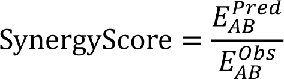

where 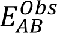 denotes the observed combined effect of drug A and B at concentration X and Y, respectively. For example, if drug A and B suppressed tumor growth 50% and 50% compared to the control group, then the predicted combined effect would be 75% (= 0.5 + 0.5 – 0.5×0.5). If the actual inhibition of tumor growth when combined is greater than 75% (e.g., for 90%, SynergyScore = 0.89), then the compounds would be synergistic. If the tumor inhibition is less than 75% (e.g., for 60%, SynergyScore = 1.25), then the compounds would be defined as antagonistic (Fig. 11b).

**Figure. 11.**
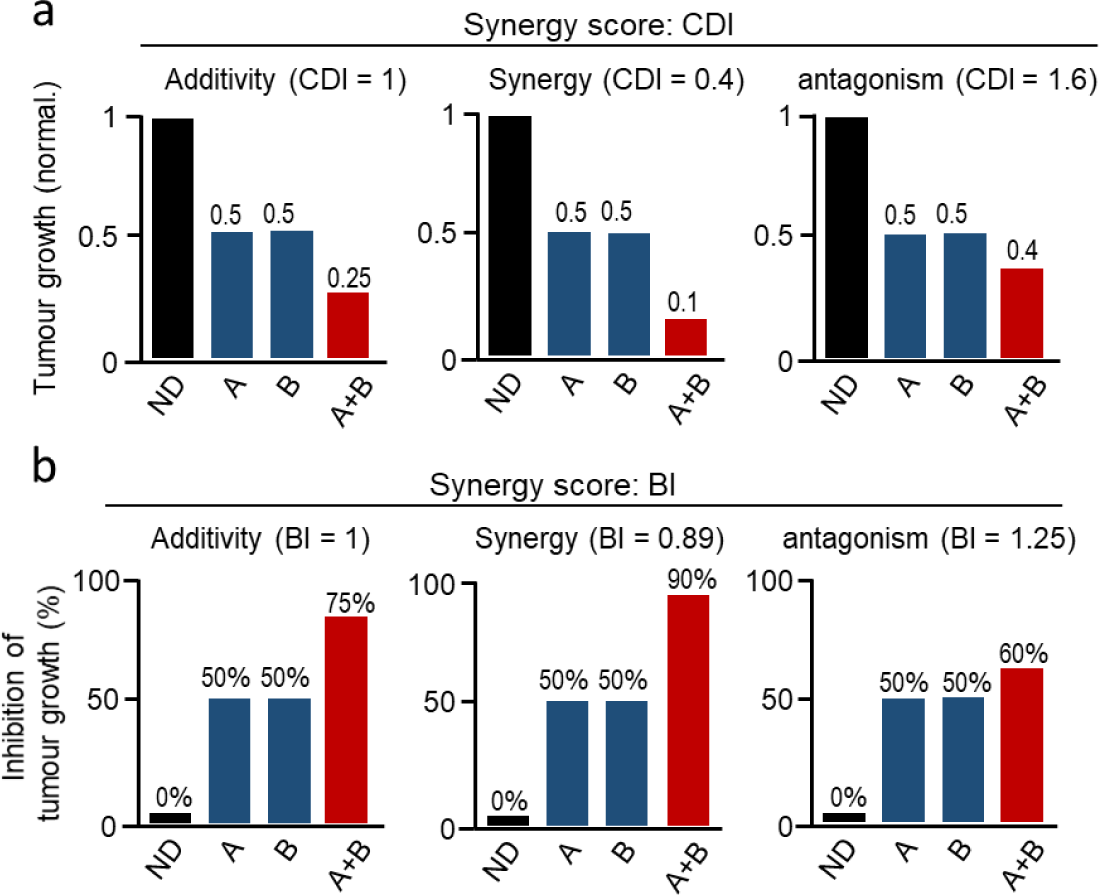
Calculation of drug synergy score. For CDI (a), drug response (e.g., tumor growth) is normalized to its untreated control. Thus, “1” indicates no drug treatment or control. “0” indicates complete inhibition. For BI (b), the drug effect indicates a percentage inhibition of the response. Thus, 0% denotes an untreated control. 100% indicates complete inhibition.

### **5.** Drug Combination Setup and Drug Synergy Tools

Because the synergy between two drugs can depend strongly on the specific doses at which they are combined, in order to gain a broad understanding of the synergy landscape it is good to perform drug combination experiments (computationally or experimentally) at various dose combinations in a “dose-matrix” setup. Shown in Figure 12, a typical set up involves combination of gradually increasing doses of two drugs A and B, around their respective IC50 values, including single-drug treatments. For example, the drugs could be combined at five doses: 1/5, 1/2, one, two and five folds of the IC50 concentration (Fig. 12). The first row and column of the matrix correspond to the single-drug treatments, and the first element of the matrix corresponds to vehicle control (UTC).

**Figure 12.**
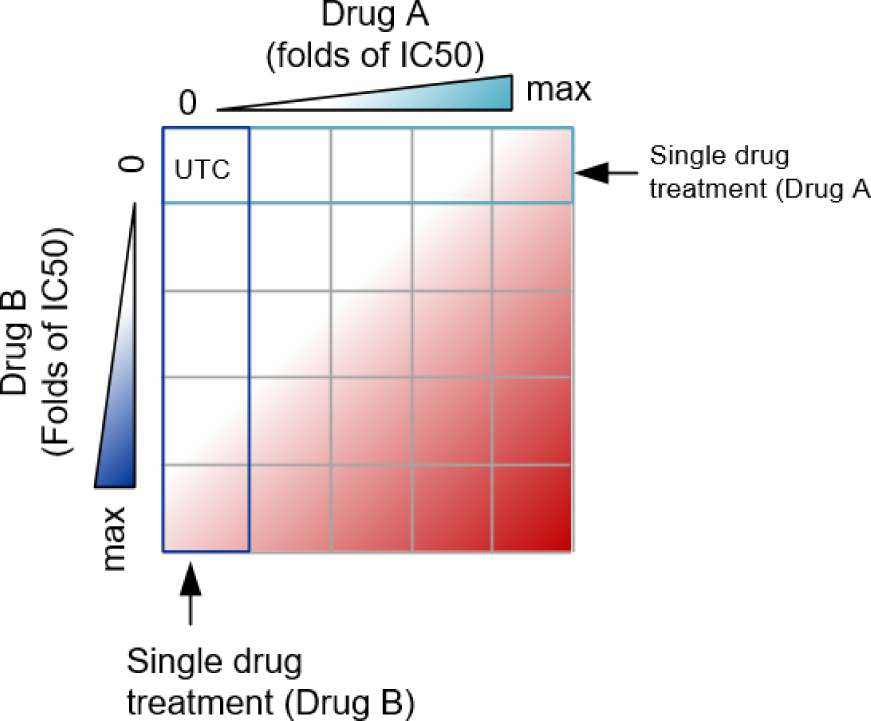
A dose-response matrix. The first row and column indicate effects of a single drug treatment. The first element of the matrix corresponds to the untreated control (UTC). max: the maximal tolerance concentration.

In addition to the CDI and BI methods for computing drug synergy, there are other methods, including Chou-Talalay (32), HSA (33), Loewe (58), and ZIP (34, 35). A number of software tools are available that allow one to compute drug synergy using the data obtained from the dose-matrix experiment setup above. For instance, SyneryFinder (35) is a web-based application that allows interactive analysis and visualization of drug synergy using a variety of methods, such as HSA (33), Loewe (58), BI (30, 31), and ZIP (34, 35). A stand-alone implementation of SyneryFinder as an R package is also available (59). Combenefit is a stand-alone software for Windows (60), which provides calculation of drug synergy using the Loewe, BI, and HSA methods. In addition, CompuSyn is another computer program to determine drug synergy using the median-effect principle of the mass-action law and the combination index theorem (32, 61) as part of the Chou-Talalay method (https://www.combosyn.com/).

## Acknowledgements

This work was supported by a Victorian Cancer Agency Mid-Career Research Fellowship (MCRF18026), an Investigator Initiated Research Scheme grant from National Breast Cancer Foundation (IIRS-20-094); and a Metcalf Venture Grant by Cancer Council Victoria, Australia awarded to L.K.N.

## Appendix A. Ordinary Differential Equations of the EGFR-MET Model

The reaction rates are given in Appendix B.

| Left-hand sides | Right-hand sides | Initial conditions (nM) |
| --- | --- | --- |
| $d[pEGFR]/dt$ | $v1 - v2$ | 0.109 |
| $d[EGFRub]/dt$ | $v3 - v4$ | 6.940 |
| $d[PYK2m]/dt$ | $v5 - v6$ | 0.622 |
| $d[PYK2]/dt$ | $v7 - v8 - v9 + v10$ | 9.299 |
| $d[pPYK2]/dt$ | $v9 - v10$ | 2.510 |
| $d[pSTAT3]/dt$ | $v11 - v12$ | 1.178 |
| $d[cMETm]/dt$ | $v13 - v14$ | 0.023 |
| $d[cMET]/dt$ | $v15 - v16 - v17 + v18$ | 4.672 |
| $d[pcMET]/dt$ | $v17 - v18$ | 0.502 |
| $d[pCbl]/dt$ | $v19 - v20$ | 10.476 |
| $d[aPTP]/dt$ | $v21 - v22$ | 0.494 |
| $d[pERK]/dt$ | $v23 - v24$ | 0.669 |
| $d[STAT3uStattic]/dt$ | $v25$ | 0.00 |

## Appendix B. Reactions and Reaction Rates of the EGFR-MET Network Model

|  | Reaction | Reaction rates |
| --- | --- | --- |
| v1 | $EGFR \rightarrow pEGFR$ | $kc1 * (EGF / (1 + Gefitinib / Ki1) + caEGF) * EGFR / (Km1 + EGFR)$ |
| v2 | $pEGFR \rightarrow EGFR$ | $(Vmax2 + kc2 * aPTP) * pEGFR / (Km2 + pEGFR)$ |
| v3 | $EGFR \rightarrow EGFRub$ | $(Vmax3 + kc3 * pCbl) * EGFR / (Km3 + EGFR) * Ki3a / (Ki3a + PYK2_{tot} / (1 + PF396 / Ki3b))$ |
| v4 | $EGFRub \rightarrow EGFR$ | $Vmax4 * EGFRub / (Km4 + EGFRub)$ |
| v5 | $\emptyset \rightarrow PYK2m$ | $Vs5 + Vmax5 * pSTAT3 / (Km5 + pSTAT3)$ |
| v6 | $PYK2m \rightarrow \emptyset$ | $kdeg6 * PYK2m$ |
| v7 | $PYK2m \rightarrow PYK2$ | $Vmax7 * PYK2m / (Km7 + PYK2m)$ |
| v8 | $PYK2 \rightarrow \emptyset$ | $kdeg8 * PYK2$ |
| v9 | $PYK2 \rightarrow pPYK2$ | $(kc9a * pEGFR + kc9b * pcMET / (1 + EMD / Ki9)) * PYK2 / (Km9 + PYK2)$ |
| v10 | $pPYK2 \rightarrow PYK2$ | $(Vmax10 + kc10 * aPTP) * pPYK2 / (Km10 + pPYK2)$ |
| v11 | $STAT3 \rightarrow pSTAT3$ | $kc11 * (pPYK2 / (1 + PF396 / Ki3b)) * STAT3 / (Km11 + STAT3)$ |
| v12 | $pSTAT3 \rightarrow STAT3$ | $(Vmax12 + kc12 * aPTP) * pSTAT3 / (Km12 + pSTAT3)$ |
| v13 | $\emptyset \rightarrow cMETm$ | $Vs13 + Vmax13 * pSTAT3 / (Km13 + pSTAT3)$ |
| v14 | $cMETm \rightarrow \emptyset$ | $kdeg14 * cMETm$ |
| v15 | $cMETm \rightarrow cMET$ | $Vmax15 * cMETm / (Km15 + cMETm)$ |
| v16 | $cMET \rightarrow \emptyset$ | $(kdeg16 + kc16 * pCbl) * cMET / (Km16 + cMET)$ |
| v17 | cMET → pcMET | $(kc17*HGF+caHGF)*cMET/(Km17+cMET)$ |
| v18 | pcMET → cMET | $V_{max18}*pcMET/(Km18+pcMET)$ |
| v19 | Cbl → pCbl | $kc19*pEGFR*Cbl/(Km19+Cbl)$ |
| v20 | pCbl → Cbl | $(V_{max20}+kc20*aPTP)*pCbl/(Km20+pCbl)$ |
| v21 | PTP → aPTP | $kc21*pEGFR*PTP/(Km21+PTP)$ |
| v22 | aPTP → PTP | $V_{max22}*aPTP/(Km22+aPTP)$ |
| v23 | ERK → pERK | $(kc23a*pcMET/(1+EMD/Ki23)+kc23b*pEGFR)*ERK/(Km23+ERK)$ |
| v24 | pERK → ERK | $V_{max24}*pERK/(Km24+pERK)$ |
| v25 | STAT3 + Stattic<br>↔ STAT3uStattic | $ka25*STAT3*Stattic - kd25*STAT3uStattic$ |

## Appendix C. Best-Fitted Parameter Values Used for Simulations

| Parameter | Value | Unit | Parameter | Value | Unit |
| --- | --- | --- | --- | --- | --- |
| kc1 | 413.0475 | min<sup>-1</sup> | Km17 | 9.817479 | nM |
| Km1 | 248.8857 | nM | Vmax18 | 0.060674 | nM min<sup>-1</sup> |
| Ki1 | 1.0000 | μM | Km18 | 9.954054 | nM |
| kc2 | 1406.048 | min<sup>-1</sup> | kdeg16 | 24.49063 | min<sup>-1</sup> |
| Km2 | 3.801894 | nM | kc16 | 1.174898 | min<sup>-1</sup> |
| Vmax3 | 0.000104 | nM min<sup>-1</sup> | Km16 | 528.4453 | nM |
| Km3 | 2.285599 | nM | kc19 | 52.72299 | min<sup>-1</sup> |
| Ki3a | 0.08356 | nM | Km19 | 13.30454 | nM |
| Ki3b | 1.0000 | nM | kc20 | 35.64511 | min<sup>-1</sup> |
| Vmax4 | 11.11732 | nM min<sup>-1</sup> | Km20 | 24.32204 | nM |
| kc3 | 10.78947 | min<sup>-1</sup> | kc21 | 0.003972 | min<sup>-1</sup> |
| Km4 | 90.78205 | nM | Km21 | 52.72299 | nM |
| Vs5 | 26.54606 | nM min<sup>-1</sup> | Vmax22 | 0.034914 | nM min<sup>-1</sup> |
| Vmax5 | 34.04082 | nM min<sup>-1</sup> | Km22 | 46.45153 | nM |
| Km5 | 4.74242 | nM | Vmax2 | 112.2018 | nM min<sup>-1</sup> |
| kdeg6 | 53.57967 | min<sup>-1</sup> | Vmax12 | 7.638358 | nM min<sup>-1</sup> |
| Vmax7 | 3.349654 | nM min<sup>-1</sup> | Vmax20 | 0.048306 | nM min<sup>-1</sup> |
| Km7 | 3.334264 | nM | kc23a | 7.03E+09 | min<sup>-1</sup> |
| kc9a | 0.463447 | min<sup>-1</sup> | kc23b | 8.43E+08 | min<sup>-1</sup> |
| kc9b | 0.988553 | min<sup>-1</sup> | Km23 | 2.831392 | nM |
| Km9 | 34.91403 | nM | Vmax24 | 4.4E+09 | nM min<sup>-1</sup> |
| Vmax10 | 0.530884 | nM min<sup>-1</sup> | Km24 | 0.156675 | nM |
| Km10 | 9.141132 | nM | kc10 | 0.006109 | min<sup>-1</sup> |
| kdeg8 | 0.056624 | min<sup>-1</sup> | Ki9 | 1.65577 | nM |
| kc11 | 0.321366 | min<sup>-1</sup> | Ki23 | 13.48963 | nM |
| Km11 | 20.6063 | nM | ka25 | 127.3503 | μM<sup>-1</sup> min<sup>-1</sup> |
| kc12 | 0.00029 | min<sup>-1</sup> | kd25 | 11.74898 | min<sup>-1</sup> |
| Km12 | 11.58777 | nM | caEGF | 0.089125 | nM |
| Vs13 | 0.093756 | nM min<sup>-1</sup> | caHGF | 0.009036 | nM |
| Vmax13 | 0.354813 | nM min<sup>-1</sup> | EGFRtot | 398.1072 | nM |
| Km13 | 38.72576 | nM | STAT3tot | 144.2115 | nM |
| kdeg14 | 4.560369 | min<sup>-1</sup> | Cbltot | 174.9847 | nM |
| Vmax15 | 91.41132 | nM min<sup>-1</sup> | PTPtot | 296.4831 | nM |
| Km15 | 6.456542 | nM | ERKtot | 166.7247 | nM |
| kc17 | 0.000811 | min<sup>-1</sup> |  |  |  |

